# Expansion of haematopoietic stem cells during development

**DOI:** 10.1101/029454

**Authors:** Ion Udroiu

## Abstract

Haematopoietic stem cells (HSCs) arise in the embryo in the aorta-gonad-mesonephros region and the placenta, afterwards colonise the liver (and spleen) and finally the bone marrow, which, from birth on, remains the sole site of haematopoiesis. In mouse, after the fetal phase of rapid HSCs expansion, the number of these stem cells continues to increase after birth in order to sustain the growing blood volume of the developing infant. The number of total circulating reticulocytes seems to indicate that this growth stops around the third week of murine life, well before the whole animal stops to grow. Around the same time, a period of abundant lymphopoiesis ends. Human HSCs seem to show similar patterns during post-natal growth and these could be useful to understand the peculiar features of paediatric leukaemogenesis.

---

The haematopoietic system continuously produces mature cells responsible for oxygen and carbon dioxide transport, immune response and haemostasis. These different kind of cells (erythrocytes, platelets, macrophages, neutrophils, eosinophils, basophils, dendritic cells, B-cells, T-cells, and NK-cells)are derived, through a process of differentiation and maturation known as haematopoiesis, from a finite and rare population called hematopoietic stem cells (HSCs). As a result of an accurate balance between self-renewing and differentiative divisions, they generate several distinct lineages while maintaining an undifferentiated population.

Haematopoiesis first raises in the yolk sac, producing nucleated primitive erythrocytes maturing in the bloodstream, followed by a second hematopoietic wave (also in the yolk sac) generating erythromyeloid progenitors which will mature into enucleated erythrocytes (Tober et al., 2007). Finally, a more complex, third wave comprising HSCs arises in the aorta-gonad-mesonephros region and the placenta.

In the erythroid lineage, reticulocytes represents the stage just before the mature erythrocyte: they have lost the nucleus but still retains the reticular network of ribosomal RNA (hence their name). Since in normal conditions their lifespan is fixed, their population present a simple kinetic, governed by a dynamic equilibrium between erythroblasts maturing into reticulocytes and reticulocytes maturing into erythrocytes. Therefore, their absolute number is directly proportional to the HSCs pool (Dingli & Pacheco, 2006). In an allometric study, Dingli and Pacheco (2007) used the total number of circulating reticulocytes to correlate HSCs with body mass in children, finding a linear scaling during ontogenic growth. While this was useful to find a general trend, it may have hidden some phase-specific differences of biological and clinical relevance. The total number of circulating reticulocytes (*RT*) was calculated as the productof the age (mass)-specific blood volume (*BV*), red blood cell concentration per litre (*RBC*) and age-specific reticulocyte percentages (*R*). *BV* was estimated from the work of Raes et al. (2006), *R* and *RBC* were taken from Castriota-Scanderbeg et al. (1992).

Studying the growth of the total number of murine reticulocytes – and therefore of the HSCs pool – can give interesting hints to the human development.

**Figure 1** shows the curves of blood volume (data from Grünenberg, 1941), RBC count (data from Baas, 1970), percentage of reticulocytes (data from Udroiu et al., 2014) and total reticulocytes (calculated as *RT* = *BV*×*RBC*×*R*) in mouse. Blood volume curve can bedefined by the logistic function *BV* = 1.7/(1 + 10^(24.8–^*^d^*^)×0.05^), (*R^2^*=0.97), where *d* is the age in days. The curve of RBC count, instead, can be described by = 7 × (1 − *e*^-0.05×^*^d^*) + 2.66, (*R^2^*=0.92), while that of reticulocyte percentage by *R* = 40 × *e*^-0.07×^*^d^* + 3,(*R^2^*=0.97). Finally, the total reticulocytes curve can be described by the logistic function *RT* = 0.49/(1 + 10^(3.4−^*^d^*^)×0.07^), (*R^2^*=0.99).

**Figure 1.**
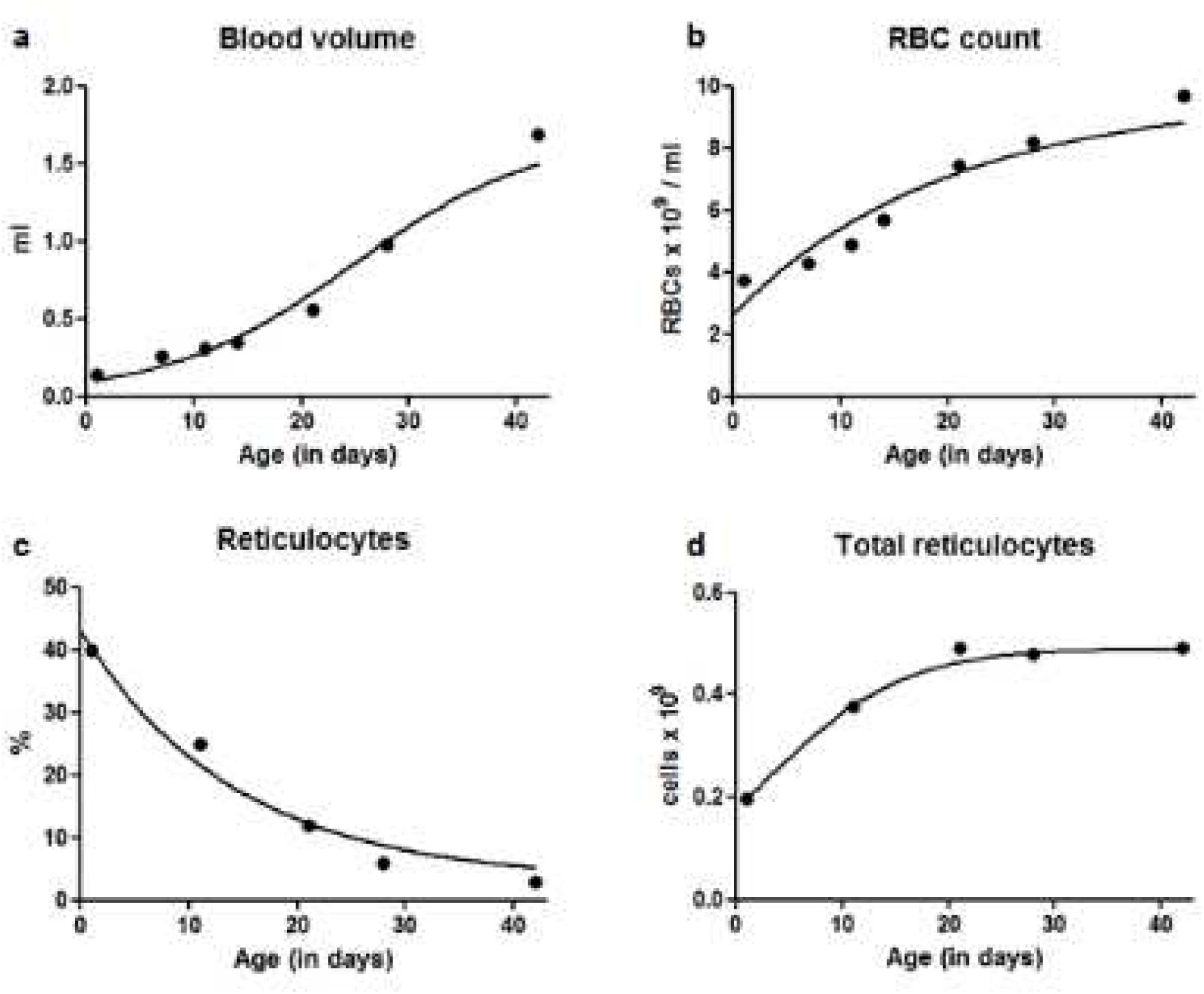
Development of bl lood parameters in mouse

Obviously these parameters may vary among strains, lab conditions etc., but our aim is not to find the exact functions for these curves. Instead, just the observation of these graphs can give useful information, especially to estimate the growth of the HSCs pool.

Blood volume shows a growth pattern strictly similar to that of the body mass of the animal, RBC count increases (with a parallel decrease of cell size) until it reaches adult values around the 6th week and the percentage of reticulocytes decreases (because of accumulation of matured erythrocytes) until it reaches a steady state at the 6th week too. These data may sound banal to a haematologist or veterinarian; however, the last graph gives an interesting clue. In fact, we can see that the total number of circulating reticulocytes reaches a plateau at the third week. This seems to be a strong evidence that three weeks after birth the HSCs pool stops to grow. Captivatingly, this agrees with findings of Bowie et al. (2007), which discovered that between the third and fourth week after birth, the HSCs pool switches from a mostly cycling population to a predominantly quiescent one. Understanding the growth dynamics of the HSCs pool may be useful for a deeper comprehension of childhood leukaemias. Whatever the model of leukaemogenesis considered (one-, two-, multiple-hits models), the fast cycling HSCs of the fetus/infant represent a bigger “target” for the leukaemogenic hits than the largely quiescent HSCs of the adult. This could explain the rapid onset of childhood leukaemias and their difference in incidence rates compared with the adult ones.

It is remarkable how the growth curve of the HSCs pool – estimated through analysis of leukocyte telomere lengths in children from birth to the age of 20 years (Sidorov et al., 2009) – resembles the growth curve of the total reticulocytes in mouse (fig. 1d). In particular, HSCs replicate ~17 times duringthe first year of life of the child, ~2.5 times/year between the ages of 3 and 13 years and then slows until reaching a rate of ~0.6 times/year in adults. This could be the reason (or one of the reasons) why paediatric leukaemias are found mostly during infancy and childhood and rarely during puberty.

Beside the switch in the self-renewing activity of HSCs, another important change takes placeat the end of infancy in mice, when the lymphoid-deficient sub-set of HSCs passes from minority to majority (Benz et al., 2012). In fact, from a HSCs pool that sustains an interleukin (IL)7-independent B-lymphopoiesis, it switches to an adult HSCs population that generates B lymphocytes in a strictly IL7-dependent manner (Kikuchi and Kondo, 2006). A similar switch in human HSCs could be related to the incidence rate of acute lymphoid leukaemia, which peaks at 1-4 years of age and then declines.

With their allometric study on total circulating reticulocytes, Dingli and Pacheco (2007) demonstrated that the number of HSCs increases during human growth. Parallelly, our calculation on murine circulating reticulocytes – in agreement with other studies (Bowie et al., 2007; Benz et al., 2012) – shows that the HSCs number stops to increase between the third and fourth week of life, i.e. before the mouse has stopped growing. During the same period, abundant lymphopoiesis also ends. Since human HSCs seem to show a similar development, further investigations are needed to: confirm the growth kinetics of HSCs during infancy, childhood and puberty; if and how the switch in lymphopoiesis takes place; how these developmental patterns are related to the specific features of paediatric leukaemias.

